# Image-based elastography of heterochromatin and euchromatin domains in the deforming cell nucleus

**DOI:** 10.1101/2020.04.17.047654

**Authors:** Soham Ghosh, Victor Crespo Cuevas, Benjamin Seelbinder, Corey P. Neu

## Abstract

Chromatin of the eukaryotic cell nucleus comprises of microscopically dense heterochromatin and loosely packed euchromatin domains, each with distinct transcriptional ability and roles in cellular mechanotransduction. While recent methods have been developed to characterize the nucleus, measurement of intranuclear mechanics remains largely unknown. Here, we describe the development of *nuclear elastography*, which combines microscopic imaging and computational modeling to quantify the relative elasticity of the heterochromatin and euchromatin domains. Using contracting murine embryonic cardiomyocytes, nuclear elastography reveals that the heterochromatin is almost four times stiffer than the euchromatin at peak deformation. The relative elasticity between the two domains changes rapidly during the active deformation of the cardiomyocyte in the normal physiological condition but progresses more slowly in cells cultured in a mechanically stiff environment, although the relative stiffness at peak deformation does not change. Further, we found that the disruption of the LINC complex in cardiomyocytes compromises the intranuclear elasticity distribution resulting in elastically similar heterochromatin and euchromatin. These results provide insight into the elastography dynamics of heterochromatin and euchromatin domains, and provide a non-invasive framework to further investigate the mechanobiological function of subcellular and subnuclear domains limited only by the spatiotemporal resolution of the image acquisition method.

## INTRODUCTION

The eukaryotic cell nucleus contains thousands of genes and long non-coding regions in the chromatin. The chromatin is compacted to fit inside a small nuclear volume through hierarchical condensation mechanisms resulting in a complex architecture^1–3^. The condensation is non-random and follows a sophisticated pattern for efficient transcription of the necessary genes. When stained, the interphase chromatin shows two distinctive regions – the heterochromatin and the euchromatin^4^. Several studies suggest that the darkly stained heterochromatin is the dense and tightly-packed form of DNA, is transcriptionally inactive, and plays a critical role in maintaining structural integrity of the nucleus^5,6^. The lightly-stained euchromatin is the loosely packed form of DNA and they are thought to be the transcriptionally active regions in nucleus^7^. In stem cells, the two chromatin regions are not distinct but in most differentiated tissue specific cells, the two regions are distinctively separated^8^. Although the function of these two regions are presently explained from the transcriptional viewpoint, the detailed functional difference between the two regions still remain elusive.

In the recent years, probing the mechanical properties of the heterochromatin and the euchromatin has gained significant interest because of the discovery of potential mechanisms that transduce mechanical forces into biological signals inside the cell nucleus – resulting into the emerging field of nuclear mechanobiology^9–11^. Mechanotransduction inside the nucleus is known to play critical roles in development, physiology, homeostasis and diseases^12,13^. Clear experimental results have been presented to show that the deformation and force in the nuclear microenvironment is directly related to the gene transcription^14–16^. To quantify and further understand the deformation characteristics of the nucleus, nuclear mechanics have been investigated for a long time^17,18^. Traditionally, the bulk nuclear deformation quantification and bulk mechanical properties of nuclei such as elastic modulus have been primary probed by research groups^19,20^ without delving into detailed intranuclear mechanical characterization – mostly due to technical limitations. Recently, new technologies^21^ have enabled us to calculate spatiotemporal displacements and generate high-resolution deformation maps in the nucleus from optical microscopy image data which enables our ability to ask and answer in depth questions regarding nuclear mechanobiology. However, as of now, no study exists that attempted to quantify intranuclear mechanical properties such as elastic modulus distribution although having this information will expand the predictive ability to quantify cell deformation by analytical and computational models. The significant effort in this line have been pursued by the intranuclear rheology^22,23^ which tracks the passive movement of fiduciary markers such as beads but such approach has a few limitations – (1) the characterization of the mechanical properties under active mechanical loading is not possible, (2) inserting beads in the nucleus is technically difficult, specifically in several primary cells and for cells *in situ* or *in vivo*, (3) bead insertion can be a highly invasive method which compromises the cell viability. Atomic force microscopy (AFM) can also potentially probe the nuclear elasticity distribution^24–27^, but that is also technically challenging specifically because – (1) the insertion of AFM probe inside the nucleus after crossing cell and tissue barrier is difficult, (2) this technique is invasive for the cell, and therefore it can significantly disrupt the cell physiology.

Optical microscopy-based^28–30^ elastography has the potential to reveal the intranuclear distribution of mechanical property in a non-invasive way. Elastography is used to solve many engineering problems where the mechanical property needs to be quantified in a minimally invasive manner. For biological problems elastography is used in different organs at the tissue scale with multiple modalities and variations such as ultrasound and magnetic resonance imaging, proving their translational/ clinical potential^31–33^ but such modalities do not have the spatial resolution to image the intranuclear details. The basic paradigm of elastography is to use the known displacement of the domain of interest and the appropriate force/ displacement boundary condition to inversely estimate the most probable elastic parameters in the domain of interest. There are several technological challenges in achieving a reliable, high resolution spatial elastography including divergence of the solution and ill-posed problem. However, simplification of the elastography problem can be achieved by discretizing the domain of interest into localized individual subdomains where the intra-subdomain elasticity can be assumed to be uniform. Using this approach, it is possible to quantify the elastic properties of two separate domains such as heterochromatin and euchromatin in the nucleus. In this paper, we report the execution of this objective of quantifying the heterochromatin and euchromatin elastic modulus using an optical microscopy-based approach. Although the ideal elastography solution could address the mechanical properties of chromatin by highest possible resolution, a two-domain elastography is the natural first step before achieving the high resolution elastography.

In the present study, we establish, validate and demonstrate the application of ‘nuclear elastography’, a technique to quantify the elasticity of the heterochromatin and the euchromatin domains of a deforming nucleus through an elaborate microscopic image analysis-based workflow. To accomplish this multi-step process, we first quantified the displacement map in the nucleus using our previously established technique ‘deformation microscopy’^21^. Next, we executed an inverse problem solution framework to iteratively calculate the elasticity of the chromatin domains by using the already computed intranuclear displacement field. As the model system we applied the technique on the nucleus of murine embryonic cardiomyocytes cultured on a PDMS substrate thus exploiting the inherent deformation behavior of beating cardiomyocytes *in vitro*. Besides, being a highly differentiated cell, when stained, the cardiomyocyte nucleus shows distinctive regions of heterochromatin and euchromatin. To rigorously validate the nuclear elastography technique, we applied known elastic moduli to heterochromatin as well as euchromatin and under this circumstance, we deformed the nucleus using experimentally similar deformation situation such as applying same displacement boundary condition on the nuclear periphery. Then, we used the baseline known elasticity values to reproduce the inversely calculated elasticity values obtained by nuclear elastography technique. After validation, we focused on applying the nuclear elastography in several physiological and pathological conditions, thus revealing new biological insights into nuclear mechanics. We showed that the dynamics of intranuclear elasticity was altered in cardiomyocytes cultured on pathologically stiff substrates compared to cardiomyocytes cultured on soft substrates that resemble native cardiac mechanical properties. Further, we showed that the intranuclear elasticity distribution was significantly compromised after global disruption of LINC complex, but not the nesprin-3 alone, in the nuclear envelope. Our data shows that the mechanical properties of the chromatin domains are optimized to enable physiological cell function and such optimization is disrupted in the pathological condition.

## RESULTS AND DISCUSSION

### Nuclear elastography provides the relative elastic modulus of the chromatin domains

Image-based nuclear elastography requires the creation of an *in silico* model of the nucleus for further finite element analysis. We established an overall workflow of creating a three-\dimensional model of the nucleus suitable for further finite element analysis in the open source software suite FEBio (Figure 1). Next, we applied an inverse problem solution framework to characterize the mechanical property of the two chromatin domains – euchromatin and heterochromatin. The displacement map obtained through registering the undeformed and the deformed images via deformation microscopy^21^ were used to inversely characterize the intranuclear mechanics. This workflow enables to apply any material model in the inverse problem solution such as hyperelastic or linear elastic material model. Because the elastic modulus is the most generic characterization of elasticity, we used material models which involves the elastic modulus. For the hyperelastic inverse solution we used the neo-Hookean model which involves the elastic modulus (*E*) and for the linear elastic model we used the Hooke’s law which also quantifies the elastic modulus (*E*), also called Young’s modulus. In either scenario, after the complete procedure we obtained the elastic modulus of the heterochromatin (*E_h_*) and the euchromatin (*E_e_*) domains. In this paper we report the ratio of *E_h_* / *E_e_*, and not the value of *E_h_* and *E_e_* separately because with a displacement boundary condition we can only obtain the relative values, not the absolute values. Therefore, all the results in this work only determines the relative measure of the elasticities of the heterochromatin vs euchromatin and should be interpreted accordingly. Calculation of the absolute values of *E_h_* and *E_e_* require a known force boundary condition, which is experimentally demanding, and not the focus of the present work. However, the present framework can be extended to compute the absolute values of chromatin domain elasticity, which might require a multiscale imaging and image analysis modalities. Besides, we discretized the nucleus only into two domains – euchromatin and heterochromatin in this work. The same framework can be further extended to calculate the elasticity at each element irrespective their location in the nucleus. However, such effort might require more sophisticated optimization algorithm and demands a high-performance computation facility. The results of two-domain elastography presented in this work is obtained using MATLAB and several open source software packages such as FEBio, GibbonCode and TetGen using only a desktop computer, where each run takes around 20 minutes. Therefore, this technique is scalable to broader use in the academic research settings.

**Figure 1.**
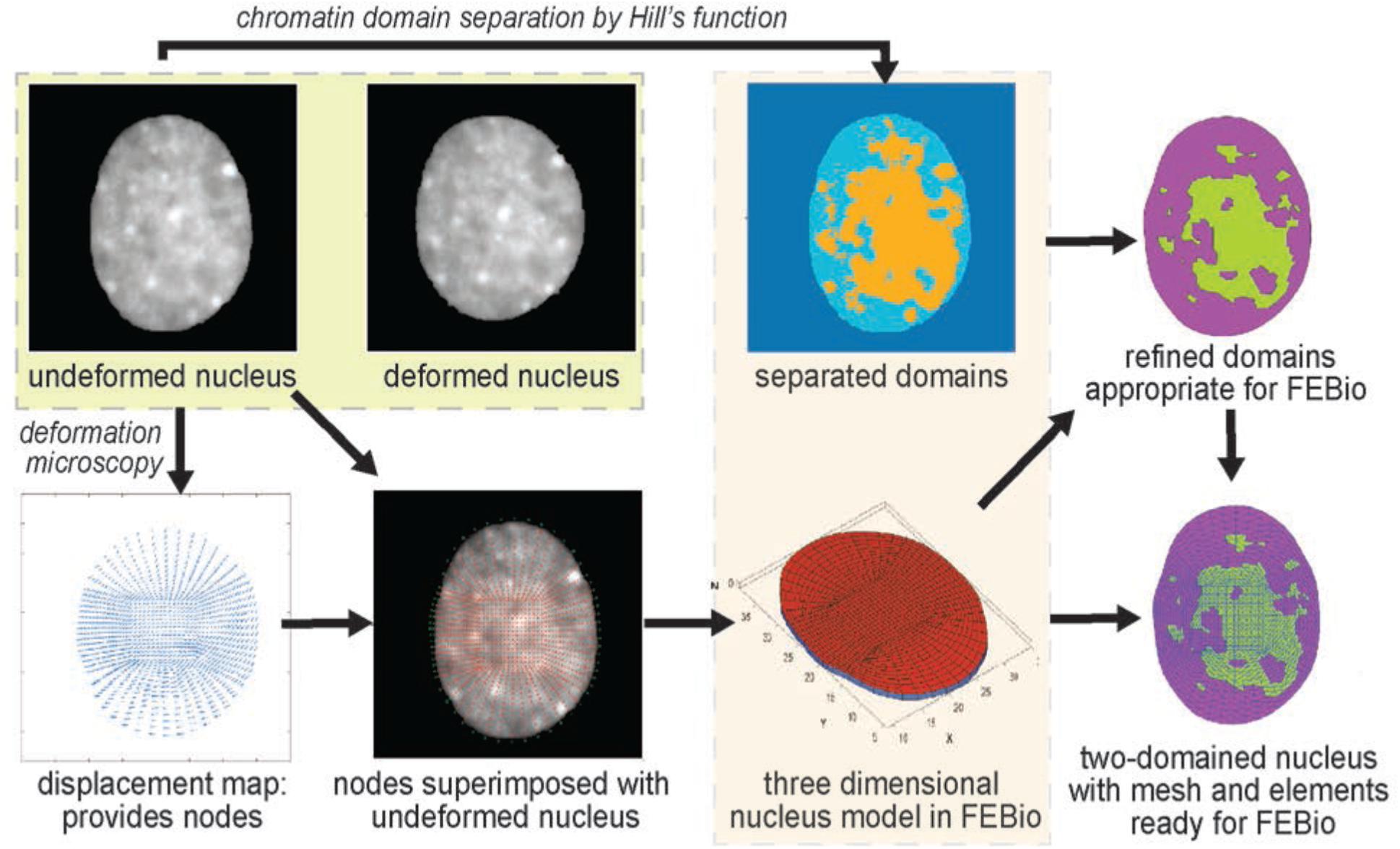
Workflow of the two-domain nuclear elastography. An undeformed image and a deformed image of the same nucleus are used to compute the relative elasticity of the heterochromatin (dense chromatin) and the euchromatin (loose chromatin) domains. First, the displacement map is obtained through deformation microscopy^21^. The heterochromatin and euchromatin domains are separated through a Hill function formulation (Figure S1 and Methods). The distribution of the nodes and their detailed information (location and connectivity) from the displacement map is used to create a three-dimensional ‘FEBio preprocessor’ model of the nucleus. Based on the separated domain map and the distribution of the nodes, the elements of the *in silico* nucleus are assigned to either the heterochromatin domain or the euchromatin domain. Further, the domains are refined to match the elements in the model nucleus, which is finally used through the FEBio interface for execution and postprocessing.

### Validation studies reveal the capability and limitations of nuclear elastography

The real experimental displacement data that we have used in this study is generated from our previous technique deformation microscopy^21^, which utilizes a new-Hookean hyperelastic material model. For the inverse solution to quantify the Elastic modulus, any material model could be incorporated. Given that we could use either hyperelastic neo-Hookean model or linear elastic Hookean model, we validated this technique for all possible scenarios such as forward hyperelastic model – inverse hyperelastic model, forward hyperelastic model – elastic model, and forward elastic model – inverse elastic model. The specific results for a forward hyperelastic material model and the inverse elastic material model results are summarized in Table 1. For different ratios of *E_hf_* and *E_ef_* (*f* denotes the forward problem), we obtain the inversely calculated ratio of *E_h_* and *E_e_*. We calculate the error in the inverse estimation as 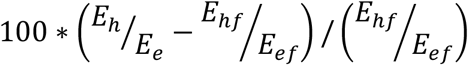. It can be noted from Table 1 that for *E_hf_* /*E_ef_* = 1, we obtain only an error of 0.41%. At a different ratio such as *E_hf_* /*E_ef_* = 0.1, we obtain an error of 9.30%. Heterochromatin is denser than euchromatin, so it is expected that heterochromatin is stiffer than euchromatin. Therefore *E_hf_* /*E_ef_* = 0.1 is probably not a physically realistic case. However, for the case *E_hf_*/*E_ef_* = 10 we obtain an error of 12.37%. For all the cases we found the inversely calculated displacement almost perfectly matches with the forward simulation displacement as denoted by *R^2^* values close to 1 (Figure 2a, 2b, 2c). Even for the cases of *E_hf_* /*E_ef_* = 0.1 and *E_hf_* /*E_ef_* = 10 we obtained *R^2^* values close to 1. This result suggests that although the optimization algorithm matches the displacement efficiently, an error occurs in the *E_h_*/*E_e_*. To understand the reason, we continued the validation study with the cases for the forward elastic material model and the inverse elastic material model (Table S1) as well as for the forward hyperelastic material model and the inverse hyperelastic material model (Table S2). In all cases, we found the error almost approaches to 0% even for extreme ratios such as *E_hf_* /*E_ef_* = 0.1 and *E_hf_* /*E_ef_* = 10. As expected, in these cases as well, the displacement from forward simulation and the inverse problem solution closely matches (Figure 2e, 2f, Figure S2). This data confirms that the error incurred in the inverse estimation as elaborated in of Table 1 is caused by the discrepancy in the material model used in the forward simulation (hyperelastic) vs inverse solution (elastic).

**Table 1.** Summary of the validation of two-domain nuclear elastography for the forward hyperelastic material model and the inverse elastic material model for a range of elasticity ratios.

| Forward problem |  |  | Inverse solution |  |  | Error |
| --- | --- | --- | --- | --- | --- | --- |
| $E_{hf}$ | $E_{ef}$ | $E_{hf}/E_{ef}$ | $E_h$ | $E_e$ | $E_h/E_e$ | |
| 1000 | 1000 | 1 | 991 | 987 | 1.004 | 0.41% |
| 1000 | 10000 | 0.1 | 276 | 2525 | 0.109 | 9.30% |
| 10000 | 1000 | 10 | 2550 | 291 | 8.763 | 12.37% |

**Figure 2.**
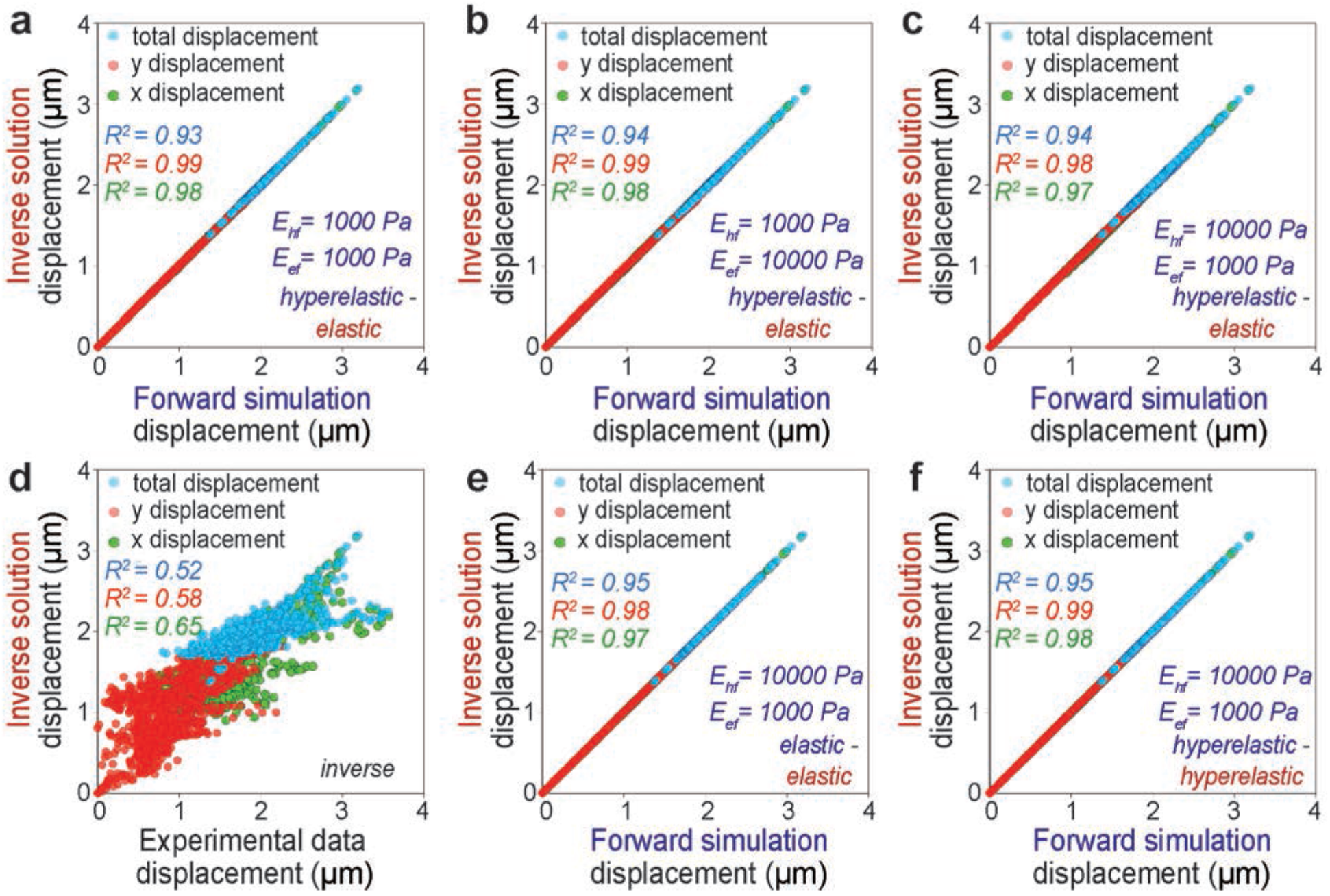
Displacement in x direction, displacement in y direction and total displacement in the xy plane is the used as the measure of the accuracy of the two-domain nuclear elastography. The upper row shows the displacements from the forward hyperelastic material model and the inverse elastic material model for a range of elasticity ratios – (a) *E_hf_*/*E_ef_* = *1:1*, (b) *E_hf_*/*E_ef_* = *1:10*, (c) *E_hf_*/*E_ef_* = *10:1*. Displacement data from the inverse problem solution is plotted against the actual displacement data fed to the inverse problem – in (d). The displacements from the forward elastic material model and the inverse elastic material model is plotted in (e) for a physically realistic value of *E_hf_*/*E_ef_* = *10:1*. The displacements from the forward hyperelastic material model and the inverse hyperelastic material model is plotted in (f) for a physically realistic value of *E_hf_*/*E_ef_* = *10:1*.

For the real experimental data, the displacement values after the completion of the inverse problem solution do not perfectly match with the input experimental displacement (Figure 2d), which will most likely cause an error in the calculation of *E_h_*/*E_e_*. To understand the reason behind this and potential impact of this error in the workflow, we continued with the next level of validation studies to assess our degree of confidence in the *E_h_*/*E_e_* value for real experimental data. We reveal that the large displacement incurred by a realistic nuclear deformation scenario is one of the responsible factors behind the mismatch between displacement calculation and resulting error in the elasticity ratio calculation (Figure 3a, 3b, 3c). Even for *E_hf_* /*E_ef_* = 10, at smaller displacement factors such as 0.05 and 0.1, which represent a small deformation, the error is almost zero (Table S3) and we obtain *E_h_*/*E_e_* = 10. With a displacement factor of 0.5, the error increases to 5.96%, and at a displacement factor = 1, we reach an error of 12.37% (Table 1). At even higher displacement factors such as 1.5, 2 and 3 the error magnifies drastically (Table S3). This data suggests that for large deformations, which is the case in real experimental data, the technique becomes less robust.

**Figure 3.**
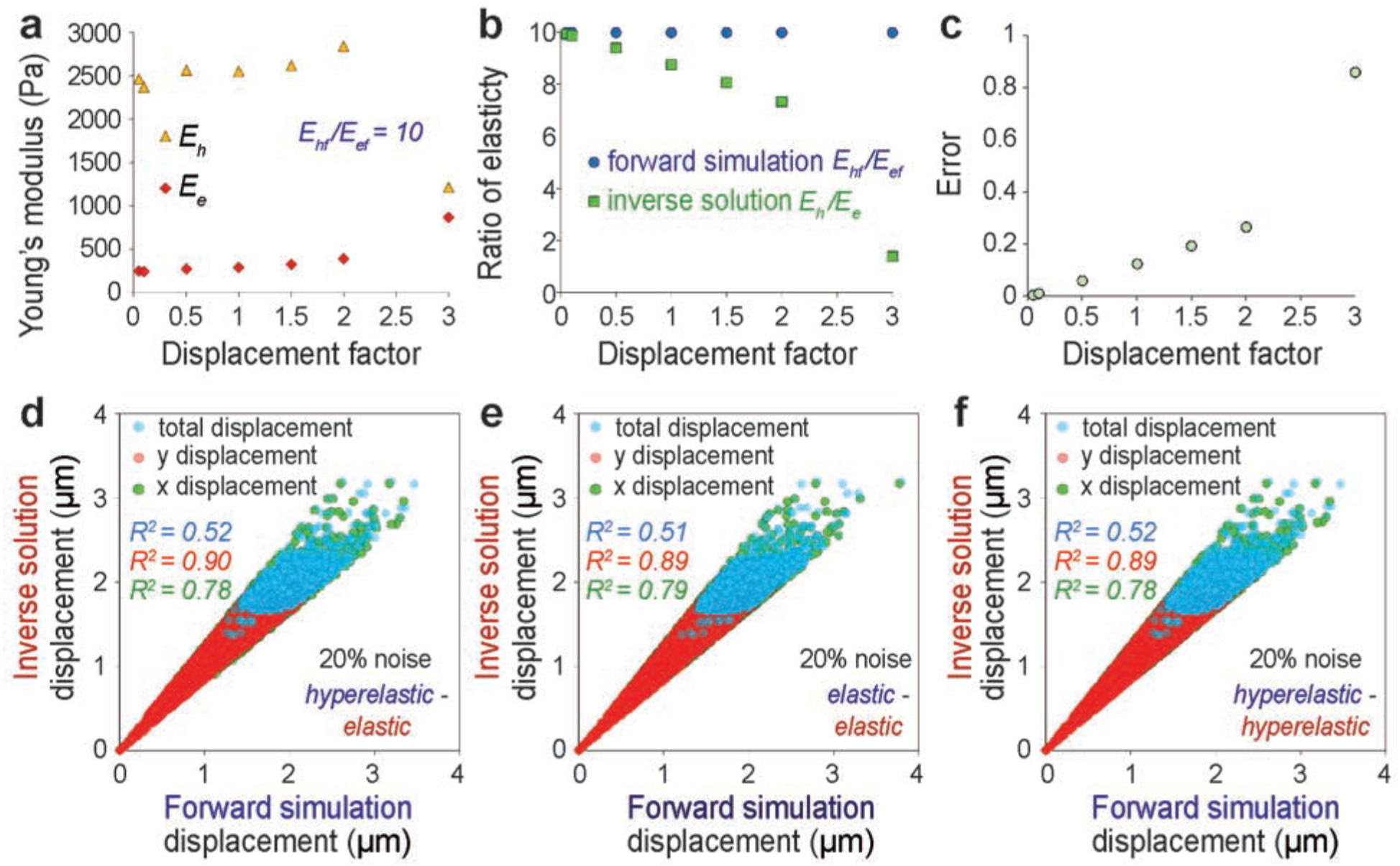
Effect of large displacement and noise on the accuracy of the elastography. For predetermined forward elasticity values of *E_hf_* = *2500 Pa* and *E_ef_* = *250 Pa*, at different displacement factors (see Methods), the *E_h_* and the *E_e_* values are inversely computed (a), their ratio is presented (b) and the error in such calculation is shown (c). Effect of 20% noise in the displacement data is investigated and the displacements are plotted for different combinations of forward model and inverse solution – (d) forward hyperelastic material model and the inverse elastic material model, (e) forward elastic material model and the inverse elastic material model, (f) forward hyperelastic material model and the inverse hyperelastic material model.

Another contributing factor that introduces error in the *E_h_*/*E_e_* calculation is the noise in the displacement data, which might be created by the inherent noise in the digital image or the error created by the displacement calculation. For every combination of forward and inverse material model it can be noted that the displacement mismatch is magnified by the noise, especially at a larger displacement (Figure 3d, 3e, 3f). Accordingly, the error in the *E_h_*/*E_e_* is incurred as computed (Table S4) – for *E_hf_* /*E_ef_* = 10, we get an error of 14.54% at the noise level of 20% for the forward hyperelastic material model and inverse elastic material model. However, it was further revealed that if we use the same material model for forward and inverse simulation (Table S5, S6), the error drastically drops even at 20% of noise. At *E_hf_* /*E_ef_* = 10, the error drops to 2.85% for the forward elastic material model and the inverse elastic material model; and the error drops to 3.03% for the forward hyperelastic material model and the inverse hyperelastic material model. Therefore, we can conclude that for this technique to be applied in the present form in order to reveal difference between experimental groups, the observed value of *E_h_*/*E_e_* should be drastically different between the groups – which is a limitation of the present study.

### Pathological condition alters the dynamics of beating cardiomyocyte nuclear mechanics

Cardiomyocytes are known to modulate their beating behavior *in vitro* based on the microenvironment stiffness. Physiological beating properties are observed on substrates that resemble native cardiac stiffness (~12 kPa), while microenvironments with increased stiffness lead to a decline in contractility^34^. The stiffening of cardiac environment is associated with various pathological conditions including cardiac hypertrophy, cardiac infarction and cardiac fibrosis^35,36^. Since a decline in cardiomyocyte contractility is associated with an abrogated nuclear strain transfer^21^, with possible downstream nuclear mechanobiological implications, we aimed to understand how the altered microenvironment mechanics affects the dynamics of intranuclear mechanics specifically the relative elasticity of the two chromatin domains.

On implementation of nuclear elastography we found that under physiological condition, where cardiomyocytes were cultured on soft substrate resembling the native heart stiffness, the nucleus deforms by a large extent and over this timeframe of large deformation at peak cardiomyocyte contraction (t_1_ and t_2_), the *E_h_*/*E_e_* value is between 2.5 and 3 (Figure 4). It should also be noted that during this timeframe, the strain values of the euchromatin region (the less stiff region) is significantly higher than the strain in the stiffer heterochromatin region (Figure S3). At later timepoints (t_3_ and t_4_) of post peak-deformation, when the nucleus comes back to its resting post-systolic state, the *E_h_*/*E_e_* value drastically increases by more than three times attaining a value of at least 8.5. Also, the nuclear strain in the post peak-contraction timepoints also significantly decreases in accordance with a lower nuclear deformation (Figure S3). The sudden change in stiffness ratio is attained rapidly only over a single timepoint change (t_2_ = 312 ms to t_3_= 468 ms). It might be possible that the stiffness difference between the heterochromatin and the euchromatin regions needs to be diminished during peak nuclear deformation to accommodate a large nuclear strain.

**Figure 4.**
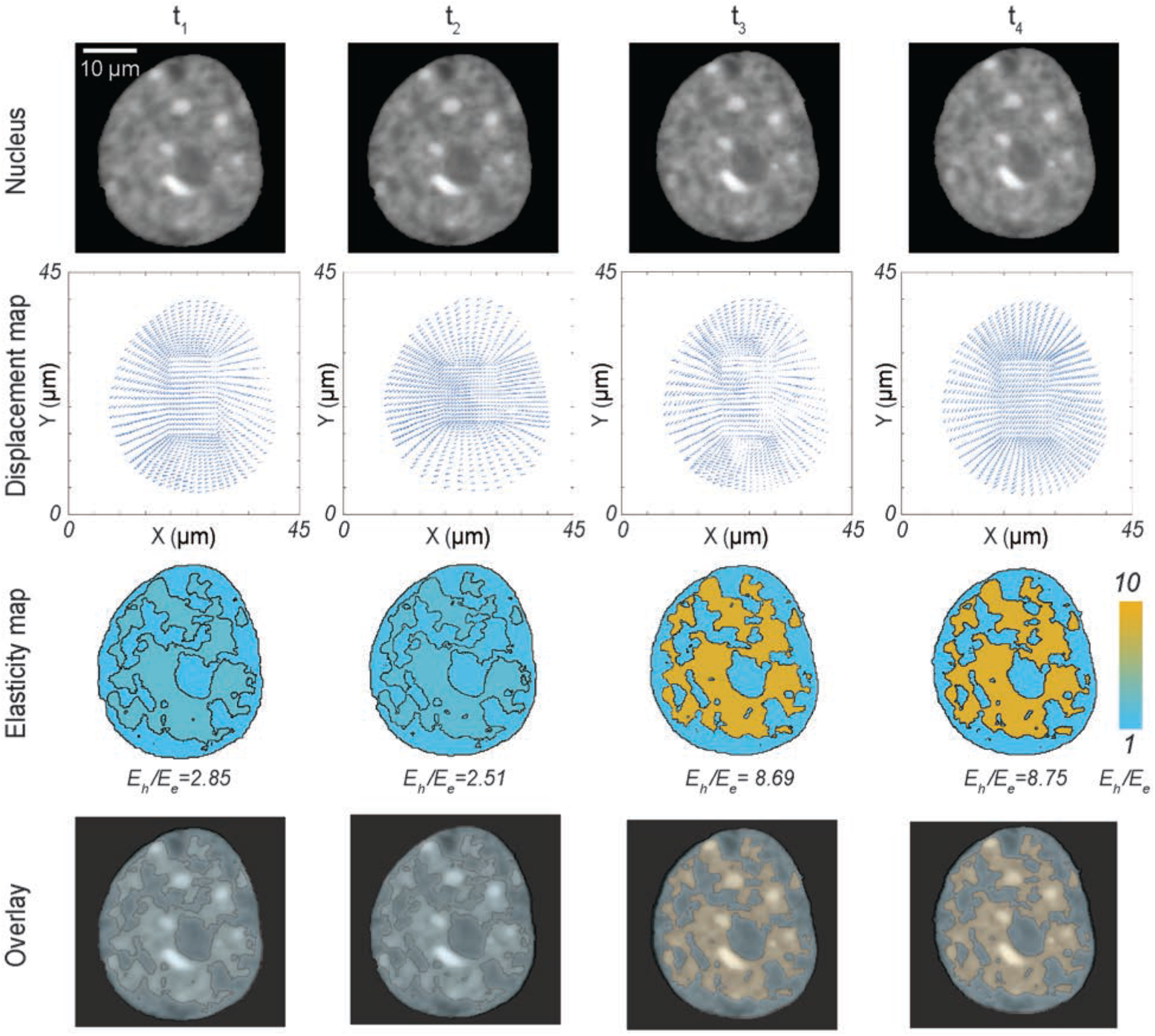
Dynamics of the relative elasticity of the heterochromatin domain and the euchromatin domain on a beating cardiomyocyte in physiological condition – with raw nuclear image, displacement map from deformation microscopy, elasticity map from two-domain elastography and the overlay of raw image + elasticity map. Timepoints for imaging and analysis are as follows: t_1_ = 156 ms, t_2_ = 312 ms, t_3_ = 468 ms, t_4_ = 624 ms.

For a stiff environment (434 kPa) which resembles the pathological cardiac condition, again we noticed a dynamic change in the *E_h_*/*E_e_* (Figure 5), however the dynamics of nuclear mechanics significantly changes from that of the soft substrate (Figure 4). The *E_h_*/*E_e_* attains a value of 5.03, when the nucleus is most deformed (Figure S4) at peak cardiomyocyte contraction. However, at a less deformed configuration (t_1_), the *E_h_*/*E_e_* value is 1.49, and at the later timepoints, where the nucleus slowly comes back to its resting state, the *E_h_*/*E_e_* value slowly adapts to 6.29 and 7.83. In essence, on stiff substrate we observe a gradual change of *E_h_*/*E_e_* over time. This observation is in stark contrast with the nucleus on soft substrate, where we observed mostly two mechanical states of *E_h_*/*E_e_*. There is a possibility that for nuclei in stiff environments, the required mechanism of abrupt change in *E_h_*/*E_e_* is compromised, which leads to an altered lower strain value (Figure S4), eventually having possible downstream mechanobiological implications.

**Figure 5.**
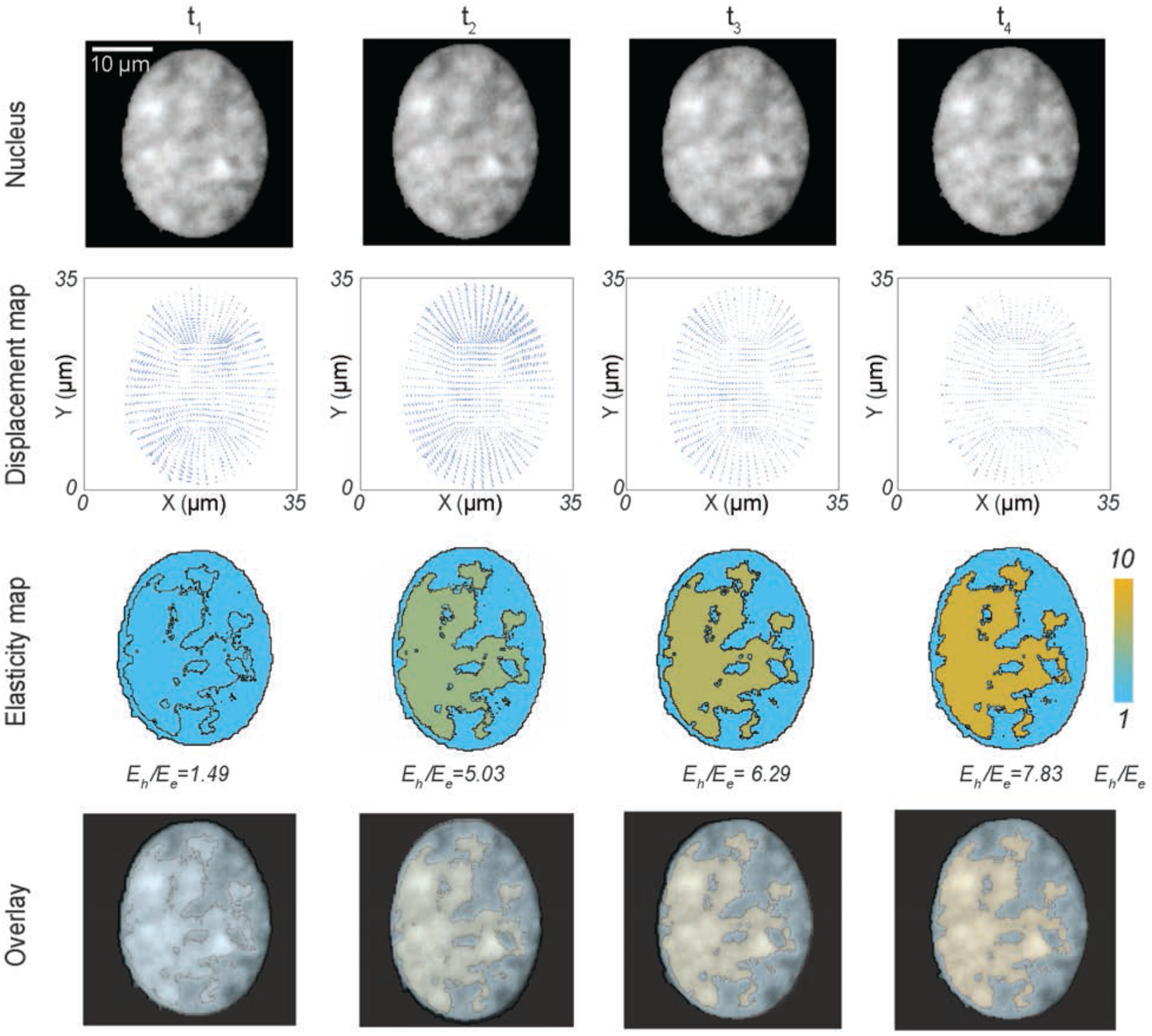
Dynamics of the relative elasticity of the heterochromatin domain and the euchromatin domain on a beating cardiomyocyte in pathological condition (stiff substrate) – with raw nuclear image, displacement map from deformation microscopy, elasticity map from two-domain elastography and the overlay of raw image + elasticity map. Timepoints for imaging and analysis are as follows: t_1_ = 156 ms, t_2_ = 312 ms, t_3_ = 468 ms, t_4_ = 624 ms.

### KASH disruption, but not nesprin-3 knock-down, affects the relative chromatin stiffness

The chromatin architecture inside the nucleus is connected to the cytoskeleton through the LINC complexes. Mutations in the LINC complexes are linked to pathological conditions^37,38^. As a second application of nuclear elastography in a cell physiology relevant problem, we hypothesized that during cardiomyocyte contraction intranuclear mechanics is compromised after knock-down of nesprin-3 or overall disruption of the LINC complex. To test this hypothesis, cardiomyocytes were transduced with vectors to knock-down nesprin-3 (shSyne3) or that overexpress a dominant-negative KASH construct (TmKash3) which displaces native nesprins via the KASH domain but lacks the cytoskeletal binding domains. As a result of these treatments we found that at peak nuclear deformation, the LINC disruption led to a *E_h_*/*E_e_* value close to 1, but the knock-down of nesprin-3 led to a value of *E_h_*/*E_e_* = 5.07 (Figure 6a). Interestingly, the overall intranuclear strain decreases for both LINC complex manipulations compared to untreated cardiomyocytes^21,39^ (Figure S5). However, while a significant change in the *E_h_*/*E_e_* value occurs after LINC disruption via TmKash3, knock down of nesprin-3 alone did not change the *E_h_*/*E_e_* value significantly at the peak nuclear deformation (Figure 6b). The stiffened microenvironments also did not significantly change the *E_h_*/*E_e_* value at peak nuclear deformation. These results suggest that the effect of nesprin-3 in determining the relative heterochromatin vs euchromatin mechanical stiffness is not significant, whereas overall LINC disruption has significantly abrupt effect on the intranuclear mechanics. We could further observe that after LINC disruption the intranuclear strain distribution is more random irrespective of the euchromatin or heterochromatin location, whereas nesprin-3 knock down showed that the higher strain is still associated with the euchromatin region, similar untreated cells on soft substrates (Figure S5). Because the chromatin architecture is connected to the nuclear envelope and through several linker molecules^12,40^, a complete disruption of KASH domain most likely isolates the chromatin architecture significantly from the nuclear envelope, thus diminishing the structural difference between euchromatin and heterochromatin, hence poising them to be mechanically similar.

**Figure 6.**
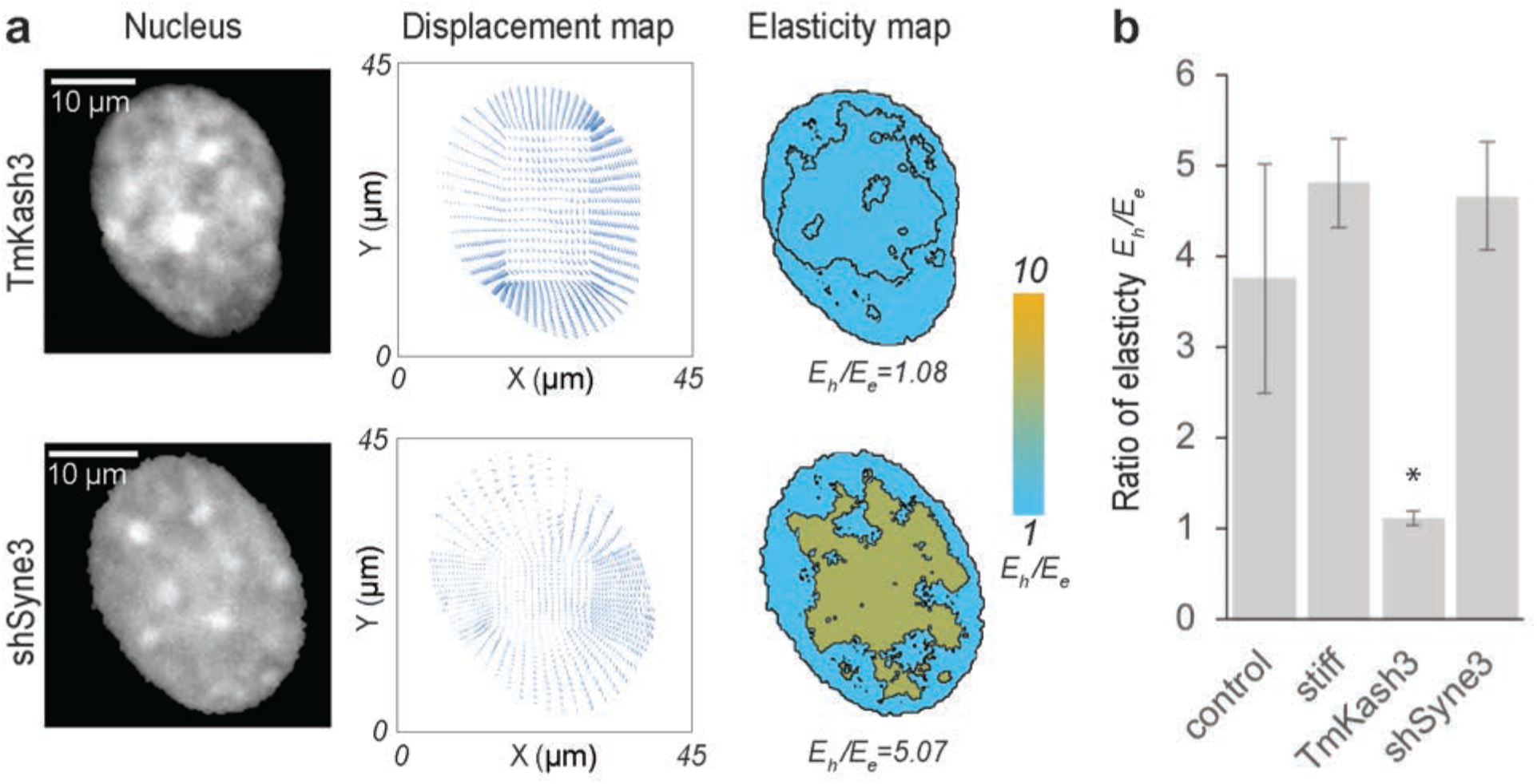
(a) Raw image of the nucleus, displacement map from deformation microscopy and elasticity map for KASH domain disrupted and nesprin-3 disrupted nuclei of beating cardiomyocytes. (b) The ratio of heterochromatin elasticity and euchromatin elasticity is shown for cardiomyocytes on soft substrates (physiological condition), stiff substrates (pathological condition), LINC complex disruption via TmKash3 and nesprin-3 knock down via shRNA interference (n ≥ 4, *p < 0.01) for a timepoint where the deformation of the nucleus was highest in each individual case (peak contraction).

### Nuclear elastography can be generalized and depends on image acquisition parameters

In this paper, we demonstrated that the relative elasticity of the heterochromatin and the euchromatin can be quantified in a non-invasive manner solely based on the images of a deforming nucleus. Previous studies indicated the possibility that heterochromatin might be stiffer than euchromatin by treating nucleus with histone methylase inhibitors which increased heterochromatin level and overall nuclear stiffness^41^, but direct proof of such possibility has not been reported so far. To the best of our knowledge the present study is the first report which mechanically delineates the euchromatin and heterochromatin using an experimental approach. The basic requirement to apply this technique is to distinguish the euchromatin and the heterochromatin from spatial image intensity, as well as obtaining a reliable intranuclear displacement map – both aspects improve with higher resolution in the microscopy images. With the rapid advent in the microscopy technology, it is possible to apply this technique in other mechanobiological problems where the intranuclear mechanics needs to be probed. Although we applied the technique in nuclei that has an endogenous green fluorescent tag, this technique can be also applied to other live nuclear imaging modalities such as Hoechst and DRAQ5^21^.

One key limitation of this technique lies in the assumption that the euchromatin and the heterochromatin regions are distinguished solely based on the image pixel intensity, which might be an oversimplification. Although the traditional biological definition of heterochromatin and euchromatin relies on the image pixel intensity in the stained nucleus, debates still exist about distinguishing the euchromatin and heterochromatin using such a simplified methodology^4^. Also, some of the low intensity pixels might be attributed to other specific nuclear bodies such as nucleolus and not euchromatin. Besides, the elasticity values we obtain from each domain is an average of the corresponding domain. A computationally demanding all element elastography can offset such assumptions and limitations, in addition to the acquisition of images with improved (e.g. super) resolution in spatial and temporal domains. which can be addressed in a future work.

## MATERIALS AND METHODS

### Cell culture

We obtained B6.Cg-Tg (HIST1H2BB/EGFP) 1Pa/J mice from Jackson Laboratory (#006069). Nuclei of the cells from homozygous mice bred from this colony display a strong fluorescence at 488 nm caused by the green fluorescence tag attached to its histone H2B. Cardiomyocytes were isolated from embryonic mice hearts 18.5 days post conception using by incubation of the tissue in 0.125% trypsin/EDTA overnight followed by 10 min digestion in residual trypsin under application of 37°C warm medium. After isolation, cells were cultured on PDMS substrates. Cardiomyocytes were cultured in Advanced DMEM/F12 containing 10% FBS at 37°C with 5% CO_2_ until the imaging experiment was carried out.

### Soft and stiff substrate

To mimic the physiological (soft) and pathological (stiff) conditions, Polydimethylsiloxane (PDMS) substrates with different formulations were used. For making soft substrate, we used Sylgard®527 ratio 1:1 (*E*=11.7 ± 5.4 kPa) and for making stiff substrate, we used Sylgard®184 (Dow Corning) ratio 1:10 (*E*=434.3 ± 54.4 kPa). Mechanical properties of the substrates were characterized as reported in our previous work^21^. For live cell imaging with 100× objective, thin (~80 µm) PDMS films were deposited on glass slides, degassed under vacuum for 30 min, cured for 2h at 100°C, and mounted on custom made cell culture dishes. Further, PDMS was ozone-treated and coated with Matrigel (Geltrex, ThermoFisher) for 1h at 37°C.

### Disruption of LINC complex

Cardiomyocytes were transduced with vectors to knock down nesprin-3 (shSyne3) or express a dominant negative KASH construct (TmKash3) that displaces native nesprins via the KASH-domain but lacks cytoskeleton binding domains. The transduction was thoroughly validated as reported in our previous publication^21^ which includes the use of a control construct that was identical to TmKash3 but lacked the KASH domain to compete with nesprins for SUN connections.

### Imaging of deforming cells

Image stacks of cardiomyocyte nuclei during contractions were captured using an inverted epi-fluorescence microscope (Nikon Ti-Eclipse) with a 100× objective and an EMCCD camera (Andor). Images were captured at 6.4 fps over a period of 10 s to visualize the entire contraction cycle of cardiomyocytes, For each nucleus, one contraction cycle was selected from the image stack to perform nuclear deformation analysis. Images of the nuclei in a post-diastolic resting state were selected as undeformed reference image.

### Displacement mapping

From the undeformed reference image and the deformed image, the displacement map was generated using our already established technique – ‘deformation microscopy’^21,42,43^. Briefly, in this technique, the undeformed image is hyperelastically warped to register with the deformed image. Deformation microscopy provides the nodal coordinate information *(x, y)* and the displacement map *d(x, y)* at each node that was further used for elastography and the strain calculations where relevant. The strains calculated were 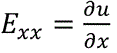 and 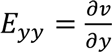, where *u* and *v* are the *x* and *y* components of the displacement *d(x, y)*. Further, hydrostatic strain is defined as the average of the E_xx_ and E_yy_: 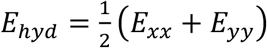.

### Two-domain elastography

The primary focus of this paper is to establish and apply the two-domain elastography to quantify the relative elasticity between the euchromatin and the heterochromatin domains of the nucleus. Therefore, this section is reported in detail. The endpoint goal of this workflow is to obtain the ratio of the heterochromatin elastic modulus (*E_h_*) and the euchromatin elastic modulus (*E_e_*).

#### Separation of chromatin domains

To distinguish the heterochromatin and the euchromatin domains in a consistent and automated manner, we applied the Hill equation^44^, a commonly used technique in biochemistry to study the binding kinetics of ligand and receptor. The rationale behind this approach is that when the pixel grayscale intensity from a raw image of nucleus is plotted after sorting by value which can range from 0 to 255, it resembles the Hill equation sigmoidal plot (Fig. S1). A sigmoidal curve was fit to the pixel grayscale intensity according to the Hill equation. The sigmoidal curve provides an inflexion point which is decided as the cut-off pixel intensity. Any pixel intensity higher than the cut-off intensity value was assigned to the heterochromatin domain, and any pixel intensity lower than the cut-off intensity value was assigned to the euchromatin domain. Therefore, a binary measure of the heterochromatin domain and the euchromatin domain was obtained.

#### Generation of nuclear model for finite element analysis framework

Using the known coordinates of the nodes obtained from deformation microscopy, we created a two-dimensional connectivity network. The 2D connectivity network was further transferred to a MATLAB-based open source Toolbox GibbonCode, which can be interfaced with the open source Finite Element Analysis package FEBio 2.6. From the 2D connectivity network, we created a 3D model consisting tetrahedral mesh using another open source code TetGen.

#### Inverse solution for elasticity calculation

To estimate the Young’s modulus of the heterochromatin (*E_h_*) and the euchromatin (*E_e_*) regions, we created an inverse problem solution framework. Briefly, we assigned two guess values (random numbers) of *E_h_* and *E_e_* in the model, applied the displacement boundary condition *d(x, y),* which is found from the experimental data, and executed the FEBio run to compute the displacement *d’(x,y)* at each node. Then, we computed the sum of root mean square error in the displacement defined as 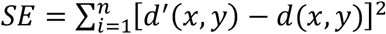, where *n* is the number of nodes. After that, we used an optimization algorithm to iteratively calculate the value of *E_h_* and *E_e_* by minimizing the objective function *SE*. For solving the optimization problem, we used our in house code based on the MATLAB function *fminsearch*. The lower and upper bounds in the function was chosen to be 100 and 2 × 10^10^ to accommodate a wide range of values for *E_h_* and *E_e_*. A plane stress condition was used in the 3D finite element model. Poisson’s ratio was assumed to be 0.35 for all cases.

#### Validation of elastography

To validate the elastography technique that was eventually applied to experimental data, we applied known material properties *E_hf_* and *E_ef_* for the two chromatin domains and executed the forward FEBio run to determine the displacement data at the nodes. Further, we used the forward run displacement data as the input *d(x, y)* to execute the inverse problem, and therefore to find the *E_h_* and *E_e_* values. We used different combinations of material model for the forward simulation and the inverse solution – (1) hyperelastic material forward model, elastic material inverse solution, (2) elastic material forward model, elastic material inverse solution, (3) hyperelastic material forward model, elastic material inverse solution. For all cases, Poisson’s ratio was assumed to be 0.35.

The hyperelastic material model utilizes the neo-Hookean model which has the parameters elastic modulus (*E*) and the Poisson’s ratio (*ν*). It employs the hyperelastic strain energy density function^45^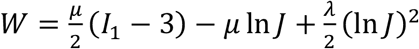, where *I_1_* is the first invariant of the right Cauchy-Green deformation tensor and *J* is the determinant of the deformation gradient tensor. μ and λ are the Lame constants defined as 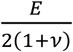 and 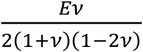 respectively. The linear elastic material model employs the Hookean model which also has the parameters elastic modulus (*E*) and the Poisson’s ratio (*ν*). However, it employs a different strain energy function 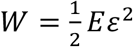, where *ɛ* is the strain.

The nuclear deformation is often associated with large displacements. Therefore, we estimated the effect of larger displacement on accuracy of the inverse problem solution. To do this, we modified the displacement solution from the forward simulation by computing *n*\* *d(x,y)*, where *n* is a constant displacement factor uniformly applied to all nodes. We applied a range of displacement factors such as 0.05, 0.1, 0.5, 1.5, 2 and 3 to accommodate 5%, 10%, 50%, 100%, 150%, 200% and 300% added displacement values to the baseline displacement *d(x,y)*.

We also estimated the effect of noise, which is inherent to the experimental data, on the accuracy of the inverse solution. For this study, artificial noise was added to the forward run displacement data as follows: *d(x,y)+d”(x,y)*, where *d(x,y)* is the mean data and *d”(x,y)* is the noise added. The noise was simulated using a uniformly distributed pseudorandom number generator and its magnitude was varied as ±5%, ±10% and ±20% of the local mean value. These magnitudes were determined from the experimental results of the present study.

### Statistics

One-way Analysis of Variance (ANOVA), followed by post hoc Tukey’s test was used to test for statistically significant differences between treatment groups. The coefficient of determination (*R^2^*) was calculated using linear regression. Error margins are reported as standard deviation (SD) about the mean, as indicated in the figure captions.

## Author contributions

Conceptualization: S.G., C.P.N.; Methodology: S.G., C.P.N.; Formal analysis: S.G., V.C.C.; Investigation: S.G., V.C.C., B.S.; Resources: C.P.N.; Data curation: S.G.; Writing – original draft: S.G.; Writing – review & editing: S.G., B.S., V.C.C., C.P.N.; Supervision: C.P.N.; Project administration: C.P.N.; Funding acquisition: C.P.N.

## Acknowledgements

This work was funded by grants from the National Science Foundation (CAREER 1349735) and the National Institutes of Health (R01 AR063712).

## Supplementary information

Supplementary Figures and Tables are available online

## Supplementary Information

**Figure S1.**
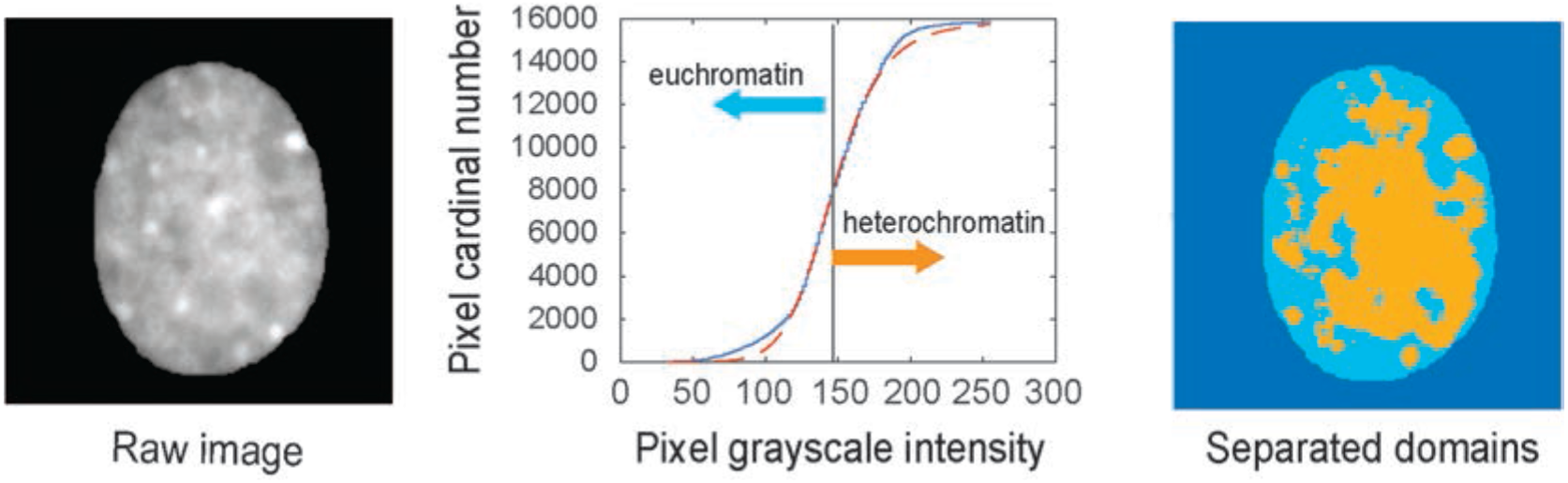
*Supporting figure for Figure 1*. The pixels of the raw image are sorted based on the grayscale pixel intensity, which can have any integer value between 0 and 255, where 0 is absolute black and 255 is absolute white. The sorted pixels are represented in the blue solid curve in the plot. Hill function fits the blue solid curve with an appropriate mathematical equation, represented by the red dotted curve. The inflexion point in the red dotted curve represents a cut-off pixel intensity. Any pixel intensity higher than the cut-off intensity value is assigned to the heterochromatin domain, and any pixel intensity lower than this cut-off intensity value is assigned to the euchromatin domain.

**Figure S2.**
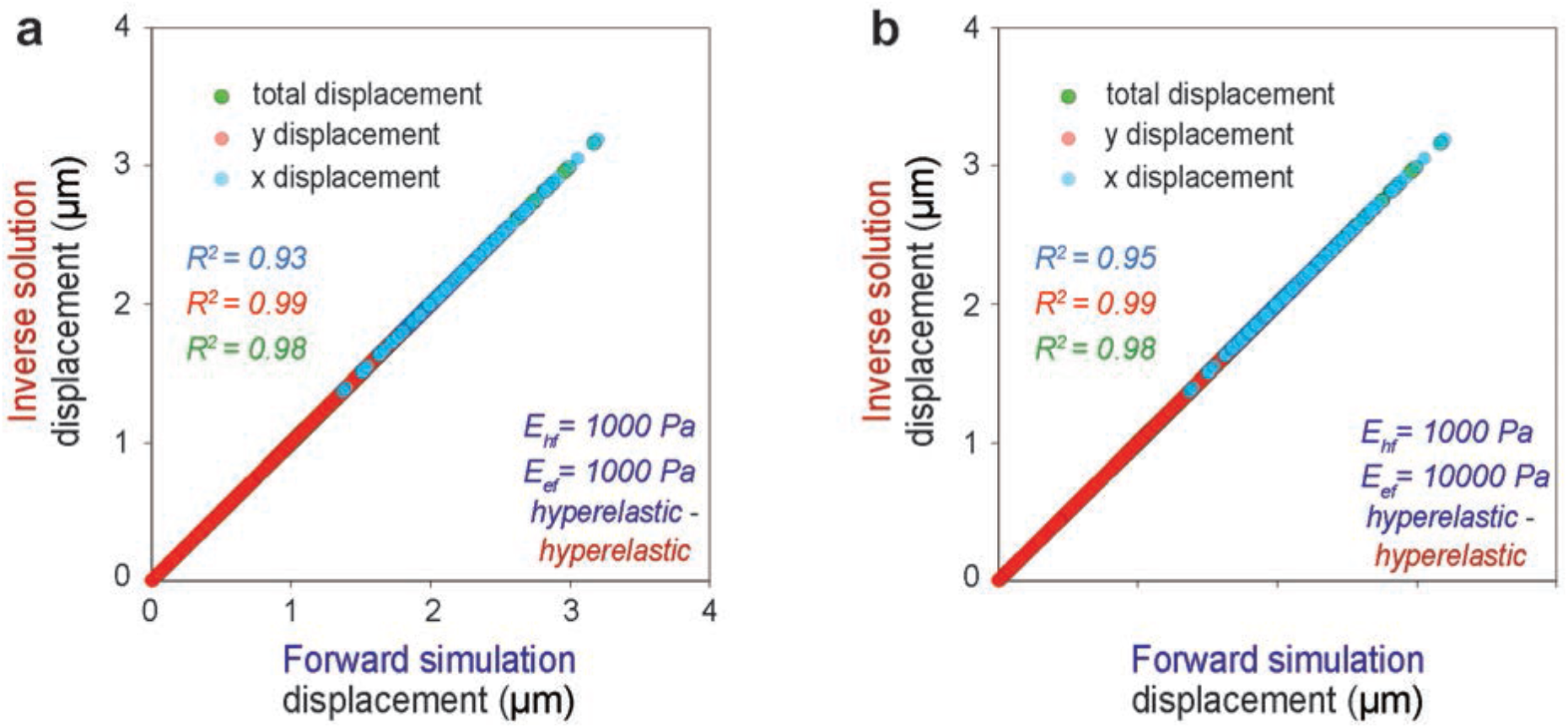
*Supporting figure for Figure 2 (f)*. Displacements from the forward hyperelastic material model and the inverse hyperelastic material model for a wider range of elasticity ratios – (a) *E_hf_*/*E_ef_* = *1:1*, (b) *E_hf_*/*E_ef_* = *1:10*.

**Table S1.**
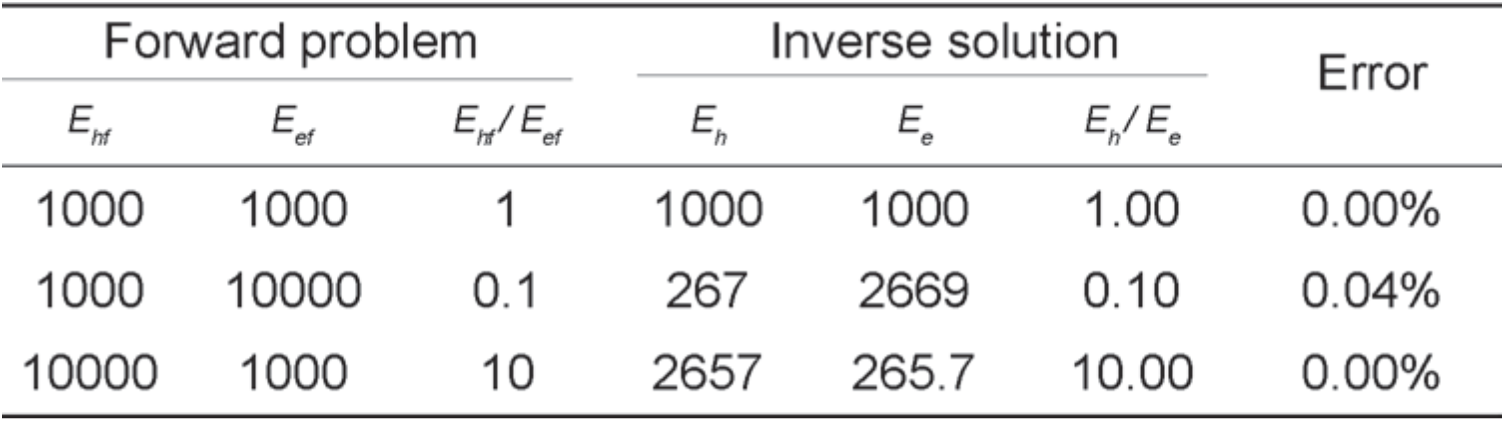
Summary of the validation of two-domain nuclear elastography for the forward (linear) elastic material model and the inverse (linear) elastic material model over a range of elasticity ratios.

| Forward problem |  |  | Inverse solution |  |  | Error |
| --- | --- | --- | --- | --- | --- | --- |
| $E_{hf}$ | $E_{ef}$ | $E_{hf}/E_{ef}$ | $E_h$ | $E_e$ | $E_h/E_e$ | |
| 1000 | 1000 | 1 | 1000 | 1000 | 1.00 | 0.00% |
| 1000 | 10000 | 0.1 | 267 | 2669 | 0.10 | 0.04% |
| 10000 | 1000 | 10 | 2657 | 265.7 | 10.00 | 0.00% |

**Table S2.** Summary of the validation of two-domain nuclear elastography for the forward hyperelastic material model and the inverse hyperelastic material model over a range of elasticity ratios.

| Forward problem |  |  | Inverse solution |  |  | Error |
| --- | --- | --- | --- | --- | --- | --- |
| $E_{hf}$ | $E_{ef}$ | $E_{hf}/E_{ef}$ | $E_h$ | $E_e$ | $E_h/E_e$ | |
| 1000 | 1000 | 1 | 1000 | 1000 | 1.00 | 0.00% |
| 1000 | 10000 | 0.1 | 267.7 | 2677 | 0.10 | 0.00% |
| 10000 | 1000 | 10 | 2642 | 264.2 | 10.00 | 0.00% |

**Table S3.**
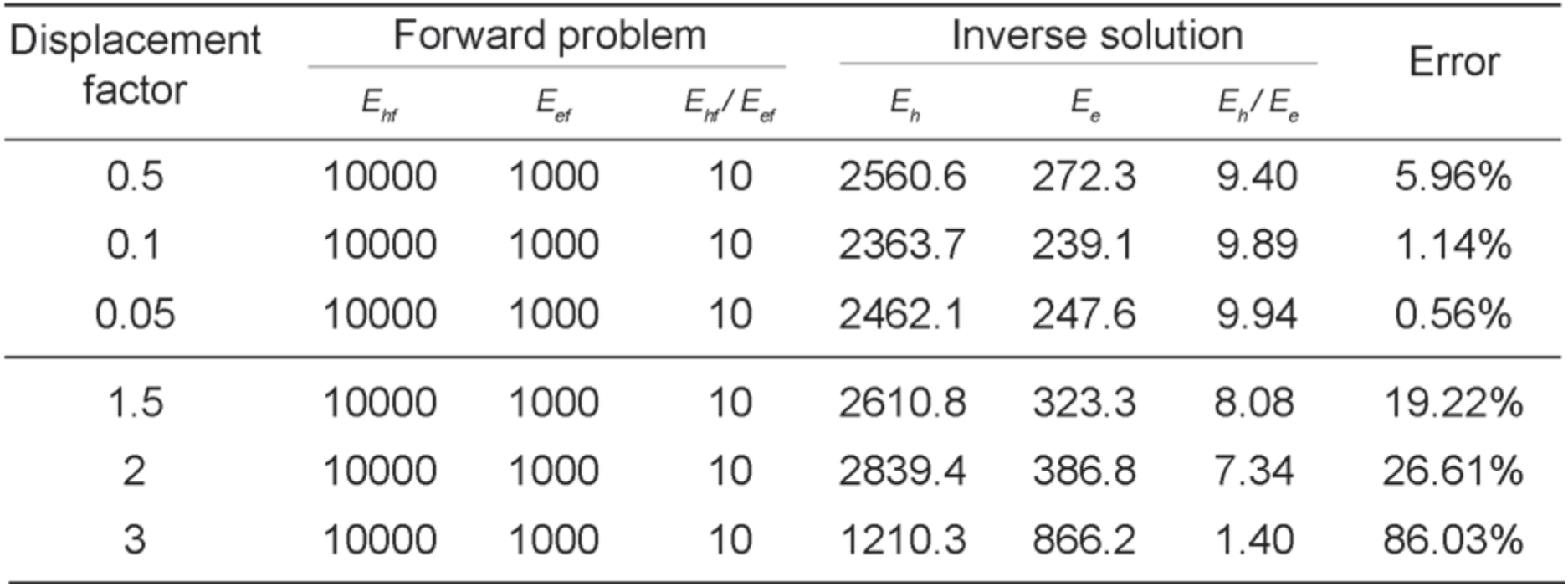
Summary of the effect of displacement factor on the elastography for the forward hyperelastic material model and the inverse (linear) elastic material model.

**Table S4.** Summary of the effect of noise on the elastography for the forward hyperelastic material model and the inverse (linear) elastic material model.

| Forward problem |  |  | Noise | Inverse solution |  |  | Error |
| --- | --- | --- | --- | --- | --- | --- | --- |
| $E_{hf}$ | $E_{ef}$ | $E_{hf}/E_{ef}$ | | $E_h$ | $E_e$ | $E_h/E_e$ | |
| 1000 | 1000 | 1 | 5% | 1075.5 | 1057.6 | 1.017 | 1.69% |
|  |  |  | 10% | 1081.8 | 1084.3 | 0.998 | 0.23% |
|  |  |  | 20% | 1149.0 | 1181.2 | 0.973 | 2.73% |
| 1000 | 10000 | 0.1 | 5% | 280.7 | 2622.1 | 0.107 | 7.05% |
|  |  |  | 10% | 309.1 | 2786.9 | 0.111 | 10.91% |
|  |  |  | 20% | 265.1 | 2428.7 | 0.109 | 9.15% |
| 10000 | 1000 | 10 | 5% | 2471.7 | 289.0 | 8.553 | 14.47% |
|  |  |  | 10% | 2913.6 | 331.7 | 8.784 | 12.16% |
|  |  |  | 20% | 2419.3 | 283.1 | 8.546 | 14.54% |

**Table S5.** Summary of the effect of noise on the elastography for the forward (linear) elastic material model and the inverse (linear) elastic material model.

| Forward problem |  |  | Noise | Inverse solution |  |  | Error |
| --- | --- | --- | --- | --- | --- | --- | --- |
| $E_{hf}$ | $E_{ef}$ | $E_{hf}/E_{ef}$ | | $E_h$ | $E_e$ | $E_h/E_e$ | |
| 1000 | 1000 | 1 | 5% | 1012.2 | 1015.2 | 0.997 | 0.30% |
|  |  |  | 10% | 1327.5 | 1342.5 | 0.989 | 1.12% |
|  |  |  | 20% | 1099.3 | 1080.2 | 1.018 | 1.77% |
| 1000 | 10000 | 0.1 | 5% | 238.0 | 2368.3 | 0.100 | 0.49% |
|  |  |  | 10% | 243.5 | 2501.5 | 0.097 | 2.66% |
|  |  |  | 20% | 259.5 | 2453.7 | 0.106 | 5.76% |
| 10000 | 1000 | 10 | 5% | 2813.9 | 279.9 | 10.053 | 0.53% |
|  |  |  | 10% | 2576.2 | 262.2 | 9.825 | 1.75% |
|  |  |  | 20% | 2359.3 | 229.4 | 10.285 | 2.85% |

**Table S6.** Summary of the effect of noise on the elastography for the forward hyperelastic material model and the inverse hyperelastic material model.

| Forward problem |  |  | Noise | Inverse solution |  |  | Error |
| --- | --- | --- | --- | --- | --- | --- | --- |
| $E_{hf}$ | $E_{ef}$ | $E_{hf}/E_{ef}$ | | $E_h$ | $E_e$ | $E_h/E_e$ | |
| 1000 | 1000 | 1 | 5% | 1133.5 | 1141.1 | 0.993 | 0.67% |
|  |  |  | 10% | 1116.2 | 1142.4 | 0.977 | 2.29% |
|  |  |  | 20% | 1107.4 | 1095.6 | 1.011 | 1.08% |
| 1000 | 10000 | 0.1 | 5% | 249.7 | 2508.9 | 0.010 | 0.47% |
|  |  |  | 10% | 268.5 | 2623.1 | 0.102 | 2.36% |
|  |  |  | 20% | 252.8 | 2513.4 | 0.101 | 0.58% |
| 10000 | 1000 | 10 | 5% | 2389.4 | 239.5 | 9.977 | 0.23% |
|  |  |  | 10% | 2693.2 | 268.9 | 10.016 | 0.16% |
|  |  |  | 20% | 2516.0 | 244.2 | 10.303 | 3.03% |

**Figure S3.**
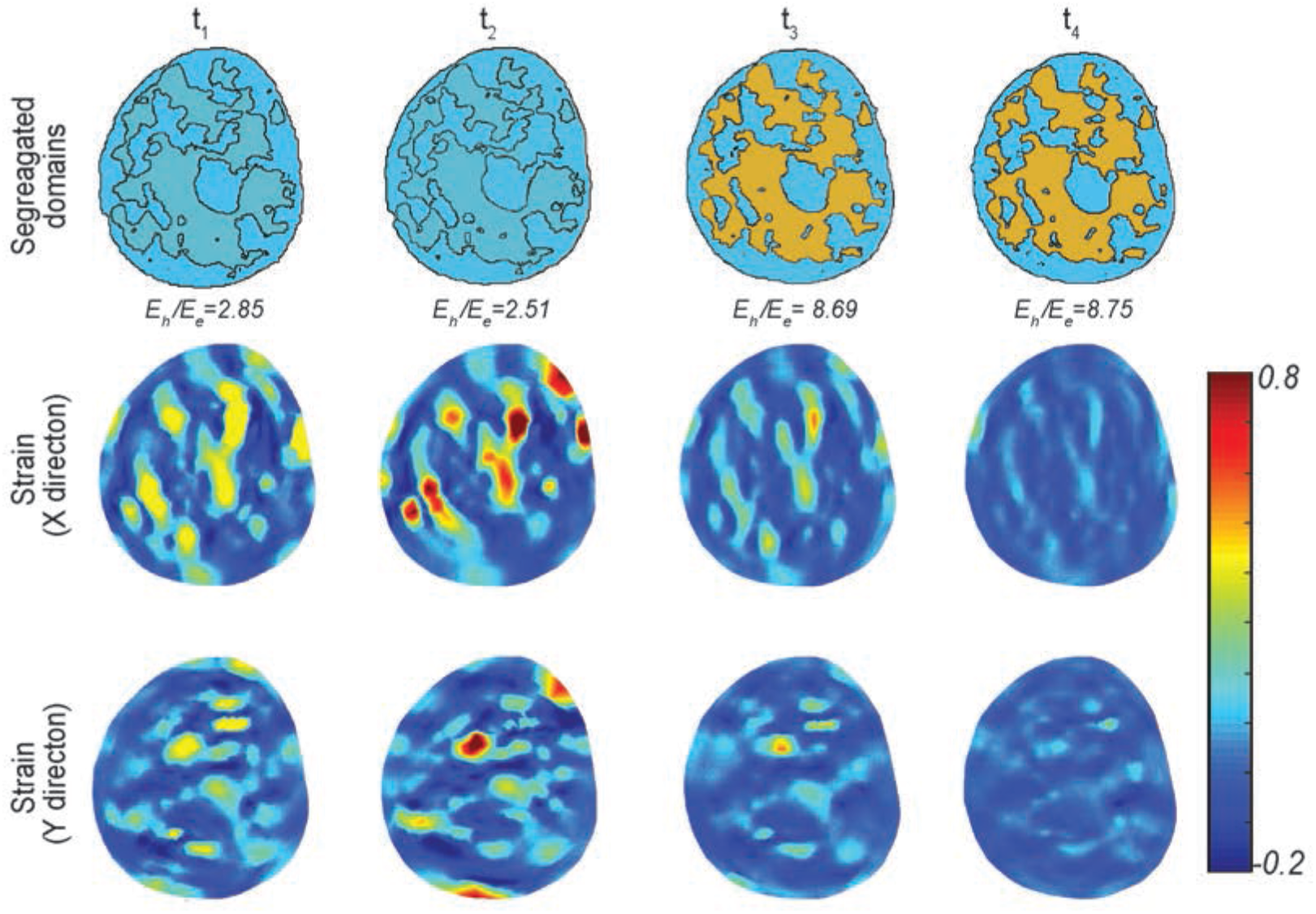
*Supporting figure for Figure 4.* Dynamics of the relative elasticity of the heterochromatin domain and the euchromatin domain during cardiomyocyte contraction plated on soft substrates (physiological condition) and its relation to the intranuclear strain map. Figure shows the elasticity map from two-domain elastography, and the strains *E*_*xx*_ (*x* directional strain), *E*_*yy*_ (*y* directional strain). Timepoints for imaging and analysis are as follows: t_1_ = 156 ms, t_2_ = 312 ms, t_3_ = 468 ms, t_4_ = 624 ms.

**Figure S4.**
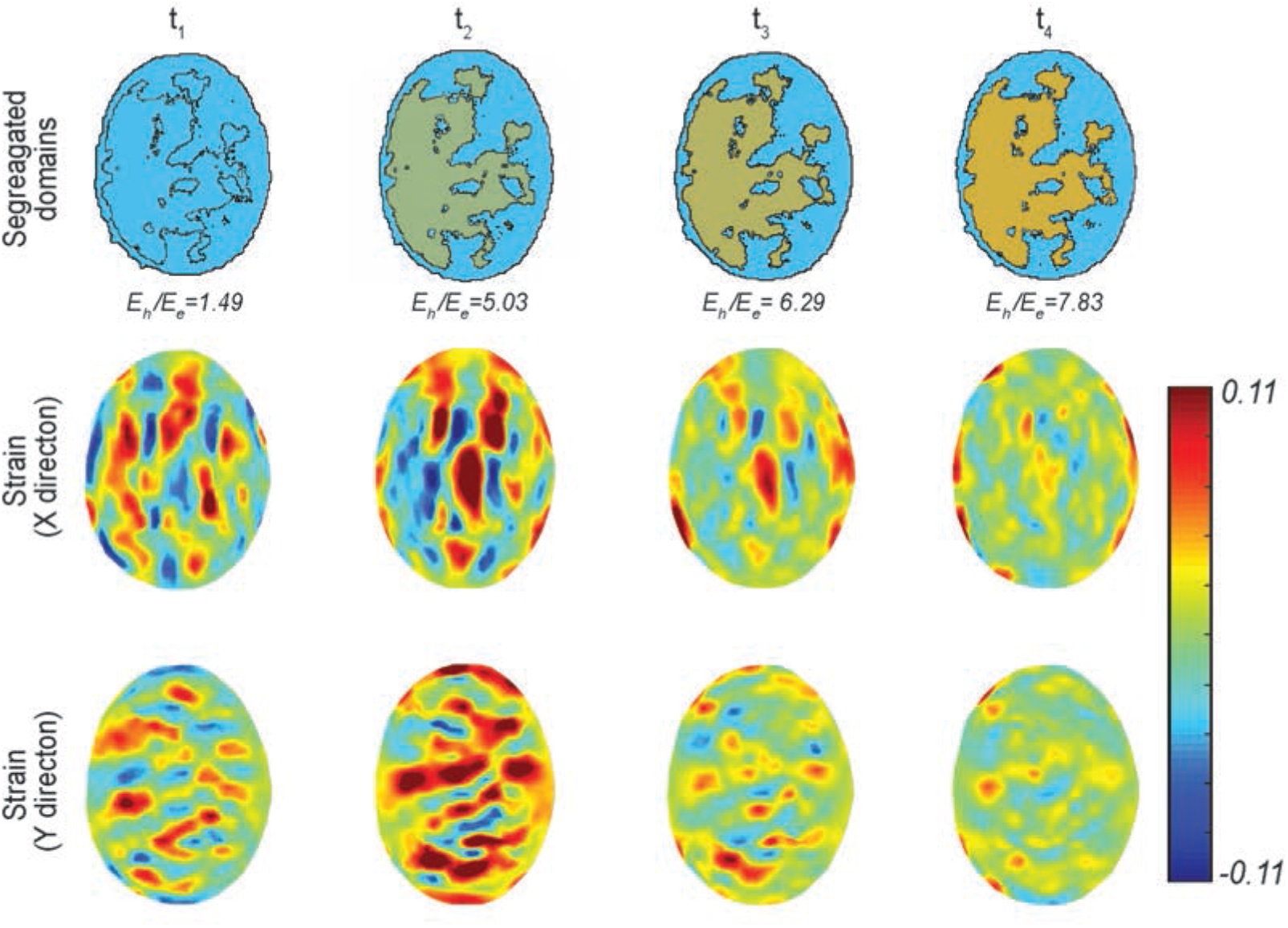
*Supporting figure for Figure 5.* Dynamics of the relative elasticity of the heterochromatin domain and the euchromatin domain during cardiomyocyte contraction plated on stiff substrates (pathological condition) and its relation to the intranuclear strain map. Figure shows the elasticity map from two-domain elastography, and the strains *E_xx_* (*x* directional strain), *E_yy_* (*y* directional strain). Timepoints for imaging and analysis are as follows: t_1_ = 156 ms, t_2_ = 312 ms, t_3_ = 468 ms, t_4_ = 624 ms.

**Figure S5.**
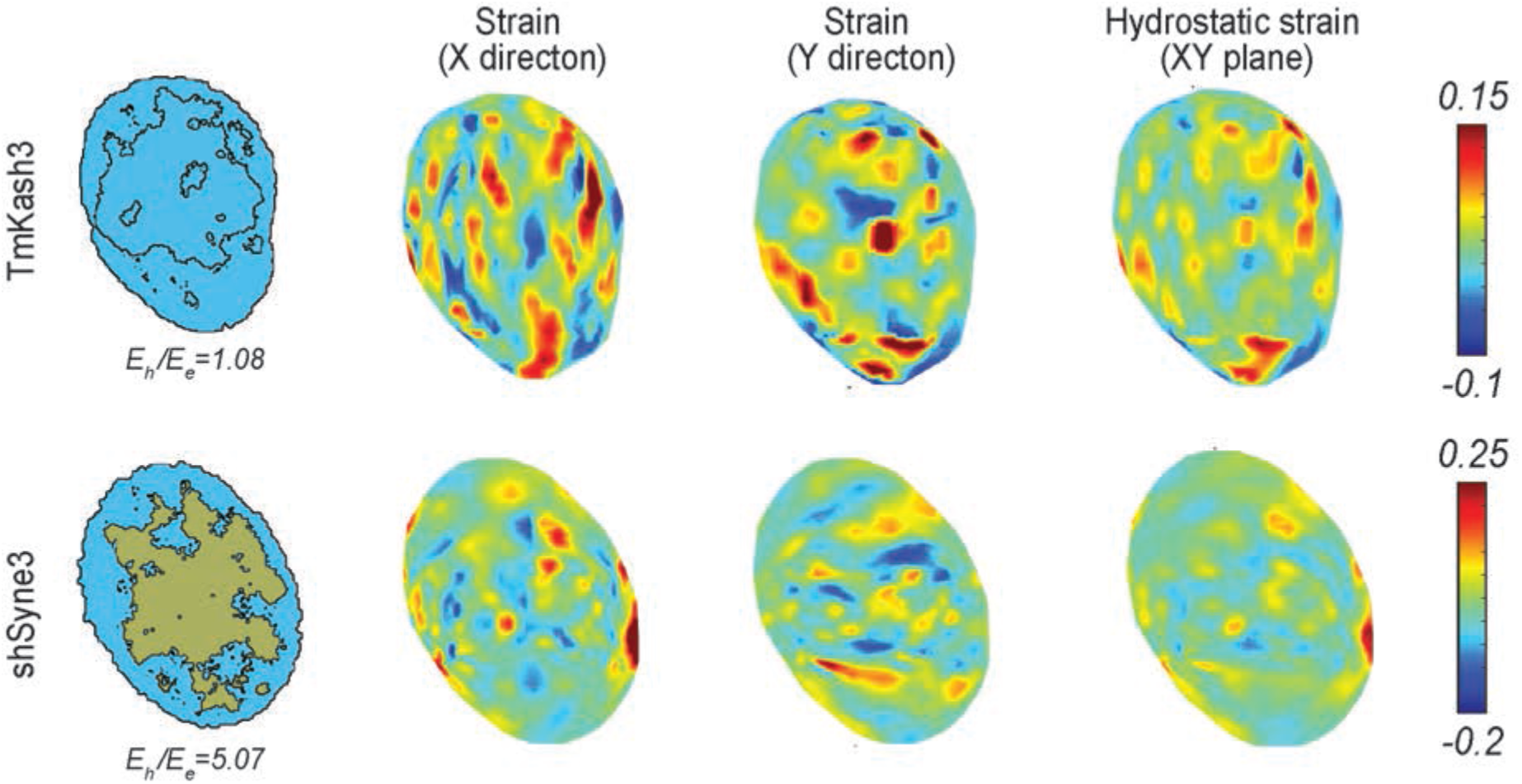
*Supporting figure for Figure 6a.* Corresponding control is represented by timepoint *t_2_* of Figure 4 and Figure S3. The figure shows the relation between the intranuclear strain map and the relative elasticity of chromatin domains after LINC disruption (TmKash3) or knock down of nesprin-3 (shSyne3). Figure shows the elasticity map from two-domain elastography, and the strains *E_xx_* (*x* directional strain), *E_yy_* (*y* directional strain), *E_hyd_* = (*E_yy_* + *E_yy_*)/*2* (hydrostatic strain in the *xy* plane).

